# Monosynaptic projections to excitatory and inhibitory preBötzinger Complex neurons

**DOI:** 10.1101/694711

**Authors:** Cindy F. Yang, Euiseok J. Kim, Edward M. Callaway, Jack L. Feldman

## Abstract

The key driver of breathing rhythm is the preBötzinger Complex (preBötC) whose activity is modulated by various categorical inputs, e.g., volitional, physiological, emotional. While the preBötC is highly interconnected with other regions of the breathing central pattern generator (bCPG) in the brainstem, there is no data about the direct projections to either excitatory and inhibitory preBötC subpopulations from other elements of the bCPG or from suprapontine regions. Using modified rabies tracing, we identified neurons throughout the brain that send monosynaptic projections to identified excitatory and inhibitory preBötC neurons. Within the brainstem, neurons from sites in the bCPG, including the contralateral preBötC, Bötzinger Complex (BötC), the nucleus of the solitary tract (NTS), parafacial region (pF_L_/pF_V_ or RTN/pFRG), and parabrachial nuclei, send direct projections to both excitatory and inhibitory preBötC neurons. Suprapontine inputs to the excitatory and inhibitory preBötC neurons include the superior colliculus, red nucleus, amygdala, hypothalamus, and cortex; these projections represent potential direct pathways for volitional, emotional, and physiological control of breathing.

## 1. Introduction

Breathing is a remarkable behavior that sustains a robust and precisely modulated rhythmic movement appropriate across all behaviors and states, including for: i) homeostatic regulation of blood gas and pH levels; ii) volitional control of airflow for speech, breathholding, etc.; iii) emotional control of airflow for laughing, crying, sighing, etc., and; iv) sleep and wakefulness. Moreover, breathing is highly sensitive to changes in stress; high levels of stress are associated with hyperventilation, while low levels of stress are associated with slow deep breathing (Boiten, Frijda, & Wientjes, 1994). The kernel of the bCPG resides in the preBötzinger Complex (preBötC), a small cluster of neurons in the ventrolateral medulla that underlies breathing rhythmogenesis to ultimately drive periodic inspiratory movements. How the brain modulates breathing by actions within the preBötC across different behaviors and states is largely unknown.

Likely substrates for such modulation are subpopulations of preBötC neurons that receive signals from regulatory brain centers such as the hypothalamus, which monitors and controls many important aspects of physiological state, e.g., temperature, arousal, and pregnancy, the limbic system related to emotional control of breathing, and cortical structures that control volitional movements that involve breathing, e.g., speech, breathhold. The regions of the brain that project to and innervate the preBötC have been roughly identified. Indeed, brain regions such as the lateral and paraventricular hypothalamus, the central amygdala, orbital cortex, and various raphe and brainstem nuclei, are sources of afferent projections to the bCPG (Gang, Sato, Kohama, & Aoki, 1995). However, retrograde tracers used in these earlier studies were not specific to neuronal subtypes, e.g., excitatory vs. inhibitory neurons, and could have been taken up by fibers of passage that could underlie false positives, motivating the present study.

Since molecular diversity across neuronal subpopulations can be a substrate for differences in afferent projections contributing to functional heterogeneity, specificity is critical when mapping a neural circuit. Particularly, the preBötC is comprised of a mix of excitatory (glutamatergic) and inhibitory (glycinergic and GABAergic) neurons that can be further segregated based on gene expression, e.g., glutamatergic excitatory preBötC neurons can express neurokinin 1 (NK1R) and/or μ-opioid receptors, the peptide somatostatin (SST), the transcription factor Dbx1 (transiently), and/or Reelin. Of these molecular subtypes, we focused on preBötC SST-expressing neurons, which are glutamatergic, and neurons that express the glycine transporter GlyT2, which comprise about half of the preBötC subpopulation of inhibitory neurons. While neither of these two populations are required for rhythmogenesis, they play a critical role in transmitting signals related to inspiratory movements to preBötC targets can modulate respiratory pattern and drive apneas, respectively (Cui et al., 2016; Sherman, Worrell, Cui, & Feldman, 2015). Determining the specific projections onto each of these subpopulations of excitatory and inhibitory neurons is critical to ultimately understanding how breathing is modulated by behavior and state.

We examined projections to excitatory SST^+^ or inhibitory GlyT2^+^ preBötC neurons by targeting modified rabies virus (EnvA+RV*dG*-mCherry) to the preBötC of Cre-expressing mice. This allows retrograde tracing restricted to one synapse, revealing neurons with direct, i.e., monosynaptic, inputs to preBötC. We found that within the brainstem, preBötC neurons receive inputs from other brainstem bCPG regions including contralateral preBötC, Bötzinger Complex, intermediate reticular region, the nucleus of the solitary tract (NTS), and parabrachial nuclei. Suprapontine regions that provide direct input to the preBötC include the periaqueductal gray (PAG), superior colliculus, substantia nigra, central amygdala (CeA), and cortex. Interestingly, neither brainstem nor suprapontine projections discriminated between excitatory or inhibitory targets, as excitatory SST^+^ and inhibitory GlyT2^+^ preBötC neurons both received projections from the same regions, suggesting that the magnitude and even sign of modulation of breathing related to emotional, cognitive, physiological, and behavioral functions may be determined by the balance of inputs to these subpopulations.

## 2. Materials and Methods

### Animals

Animal use was in accordance with the guidelines approved by the UCLA Institutional Animal Care and Use Committee. Animals were housed in a vivarium under a 12 hr light cycle with free access to food and water. All experiments were performed using adult male SST-Cre and GlyT2-Cre mice (24–30 g) that were 10–24 weeks of age at the time of AAV injection. SST-Cre mice were crossed to ROSA26-EGFP Cre-reporter mice (IMSR Cat# JAX:007914, RRID: IMSR_JAX:007914) to generate the SST reporter line used in retrograde tracing studies. Prior to surgery, animals were group housed in cages of up to 5 mice, and after surgery, animals were individually housed. GlyT2-Cre mice were kindly provided by H.U. Zeilhofer, University of Zurich, Switzerland, and all other mice were obtained from Jackson Labs (Bar Harbor, ME).

### Viruses

For rabies virus tracing, two Cre-dependent helper viruses, AAV8-EF1α-FLEX-BFP-T2A-TVA-WPRE-hGH (3.5×10^13^ GC/ml) (E.J.K./E.M.C. unpublished) (referred to in the text as AAV-FLEX-BFP-TVA) and AAV8-EF1α -FLEX -G -WPRE -hGH (7.6×10^13^ GC/ml) expressing chimeric rabies glycoprotein (PBG) (AAV-FLEX-G), were produced by the Salk Viral Vector Core. PBG is a chimeric glycoprotein consisting cytoplasmic domain of SAD B19 strain glycoprotein and extracellular domain of Pasteur virus strain glycoprotein (Kim, Jacobs, Ito-Cole, & Callaway, 2016). A 3:7 mixture of AAV-FLEX-BFP-TVA and AAV-FLEX-G was used for viral injections. EnvA-pseudotyped G-deleted rabies virus, EnvA+RV*dG*-mCherry, with the titer of 3.6×10^7^ infectious units (IU/ml), was provided by the Salk Viral Vector Core.

### Surgical procedures

All experimental procedures were approved by the Chancellor’s Animal Research Committee at the University of California, Los Angeles. For AAV iontophoresis injections, male mice (10-24 weeks old) were anesthetized with isoflurane and positioned in a stereotaxic apparatus (Kopf Instruments, Tujunga, CA). The skull was exposed with a midline scalp incision, and the stereotaxic frame was aligned at bregma using visual landmarks. A drill was placed over the skull at the coordinates corresponding to the preBötC (anteroposterior, −6.78 mm; lateral, 1.25 mm) (Paxinos & Franklin, 2004), and a hole was drilled through the skull bone to expose the cerebellum. A micropipette (~20μm tip I.D.) loaded with virus (AAV-FLEX-BFP-TVA and AAV-FLEX-PBG) was aligned at bregma (including in the z-axis), guided to the preBötC coordinates, and slowly lowered at 1 mm/min until it penetrated to a depth of −4.8 mm. A silver wire electrode was inserted into the micropipette, with the tip submerged in the viral solution, and was connected to a current source (Midgard Electronics, Watertown, MA). A current of 3μA (7 sec on, 7 sec off) was applied for 5 min, and the micropipette was left in place for 10 minutes to allow viral particles to diffuse and then withdrawn at 1 mm/min to minimize the backflow along the micropipette track. After surgery, mice were housed for 3 weeks to allow robust expression. For injection of the modified rabies virus, a second surgery was performed as above, but a Picospritzer pressure injection system was used to dispense 20-40nl of EnvA+RV*dG*-mCherry rabies (Salk Institute Viral Vector core). After 7-10 days we processed the brains for histology, using anatomical landmarks, e.g., facial and hypoglossal nuclei, and nucleus ambiguous, to verify that the starter cells were localized to the preBötC.

For anterograde tracing experiments, tetramethylrhodamine dextran (5% in saline) was injected into the central amygdala (−1.20 mm anteroposterior, ±2.40 mm lateral, −4.25 mm ventral), cortex (0.75 mm AP, ±1.25 mm lateral, −1.00 mm ventral), red nucleus (−3.40 mm AP, ±0.65 mm lateral, −3.75 mm ventral), and thalamus (−2.30 mm anteroposterior, ±1.00 mm lateral, −3.00 mm ventral). SST-GFP reporter lines were injected to identify the molecular phenotype of retrogradely labeled preBötC neurons. Iontophoresis was performed as described above for AAV injection with an application of 3μA for 7 min. After surgery, mice were house for 8-10 days before their brains were processed for histology.

### Histology

Mice transfected with AAV and modified rabies viruses were processed as previously described (Sherman et al., 2015; Yang & Feldman, 2018). Briefly, mice were deeply anesthetized with isoflurane and perfused transcardially with saline followed by 4% paraformaldehyde in phosphate-buffered saline (PBS). Brains were removed, postfixed overnight at 4°C, and embedded in 3% Bactoagar. Free-floating coronal or sagittal sections (40 μm) were collected using a vibratome (Leica Biosystems, Buffalo Grove, IL) and stored at 4°C until further processing. Sections were incubated with primary antibodies in PBS containing 0.3% Triton X-100 overnight at room temperature. After 3 washes, sections were incubated in species-appropriate secondary antibodies in PBS for 2h at room temperature. After 3 washes, sections were mounted onto gelatin-treated glass slides and coverslipped. Fluorescence was visualized with a confocal laser scanning microscope (LSM710; Carl Zeiss, Oberkochen, Germany). Images were acquired with Zen software (Carl Zeiss), exported as TIFF files, processed in Image J (NIH, Bethesda, MD; RRID: SCR_003070NIH), and assembled in Adobe Illustrator.

### Antibody Characterization

We used the following primary antibodies: rabbit polyclonal anti-somatostatin-14 (T4103, 1:500; Peninsula Laboratories, San Carlos, CA), mouse monoclonal anti-NeuN (MAB377, 1:500; Millipore, Billerica, MA), chicken polyclonal anti-GFP (GFP-1020, 1:500; Aves Labs, Tigard, OR), goat polyclonal anti-choline acetyltransferase (AB144P, 1:500, Millipore). Secondary antibodies used were donkey antirabbit rhodamine Red-X, donkey anti-chicken DyLight488, donkey anti-mouse Cy5, and donkey anti-sheep rhodamine Red-X conjugated antibodies (1:250; Jackson ImmunoResearch, West Grove, PA).

Anti-somatostatin-14 antibody (Peninsula Laboratories, Cat# T-4103.0050, RRID: AB_518614) was raised in rabbit against the first 14 aa of the synthetic peptide SST. This antibody specifically recognizes SOM-14, SOM-28, SOM-25, and [Des-Ala^1^]-SOM-14 in radioimmunoassay tests, and does not cross-react with other peptides such as substance P, amylin, glucagon, insulin, neuropeptide Y, and vasoactive intestinal peptide, or SST analogs according to the manufacturer. Preadsorption of the antibody with the SOM-14 peptide abolishes specific immunoreactivity in mouse brain sections (Scanlan, Dufourny, & Skinner, 2003).

The anti-NeuN antibody (Millipore, Cat# MAB377, RRID: AB_2298772) was raised in mice (clone A60) against the neuron-specific protein NeuN, which is expressed in most, but not all, neurons in the adult mouse brain, and not in glia (Mullen, Buck, & Smith, 1992). The specificity of this antibody for neurons has been validated using immunohistochemistry and immunoblot analysis showing antibody binding to the nuclear fraction of brain tissue, but not other organs (Mullen et al., 1992).

Anti-GFP antibody (Aves Labs, Cat# GFP-1020, RRID: AB_10000240) was raised against recombinant EGFP in chickens, and the IgY fraction containing the antibody was affinity-purified from the yolks of immunized eggs. This anti-EGFP IgY specifically recognizes the viral-mediated EGFP-expressing neurons in adult rats (Tan et al., 2008; Tan, Pagliardini, Yang, Janczewski, & Feldman, 2010). We did not observe immunolabeling in brain sections that were not infected by the virus expressing EGFP.

The anti-choline acetyltransferase (ChAT) antibody (Millipore, Cat# AB144P, RRID: AB_2079751) was raised in goat using the human placental enzyme as immunogen. Specificity of the antibody was verified by Western blotting in mouse brain lysate, where a single band corresponding to the 68-70 kDa ChAT protein was identified. In addition, immunolabeling in brain sections using this antibody was abolished after preincubation with rat recombinant choline acetyltransferase (Marquez-Ruiz, Morcuende, Navarro-Lopez Jde, & Escudero, 2007).

### Data Availability Statement

The data that support the findings of this study are available from the corresponding author upon reasonable request.

## 3. Results

### Targeting SST^+^ and GlyT2^+^ preBötC neurons

We identified sources of direct afferent input to subpopulations of preBötC neurons using transsynaptic retrograde labeling with modified rabies virus (Kim et al., 2016; Wickersham, Sullivan, & Seung, 2013). This genetic strategy requires two stages (Fig 1): i) Introduction of Cre-dependent helper viruses into the preBötC of SST-Cre or GlyT2-Cre mice for genetic specificity. The helper viruses, AAV-FLEX-BFP-TVA and AAV-FLEX-G, encode for the avian sarcoma leucosis virus glycoprotein receptor, TVA (not normally expressed in mammals), and for an optimized rabies glycoprotein required for retrograde transport, respectively. ii) Subsequent delivery of an EnvA-pseudotyped rabies virus for retrograde tracing. Since the rabies virus cannot infect cells in the absence of the TVA receptor, infection is limited to Cre^+^ cells expressing the helper AAV. This modified rabies virus bears a deletion of the endogenous glycoprotein G that is replaced with the coding sequence for mCherry (EnvA+RV*dG*-mCherry), rendering it incapable of retrograde transport. Thus, only in Cre^+^ cells complemented with the glycoprotein via helper AAV injection will the rabies virus be able to cross the synapse to label presynaptic neurons. Presynaptic neurons do not express the glycoprotein, thus restricting viral transfer and ensuring monosynaptic retrograde spread.

**Figure 1.**
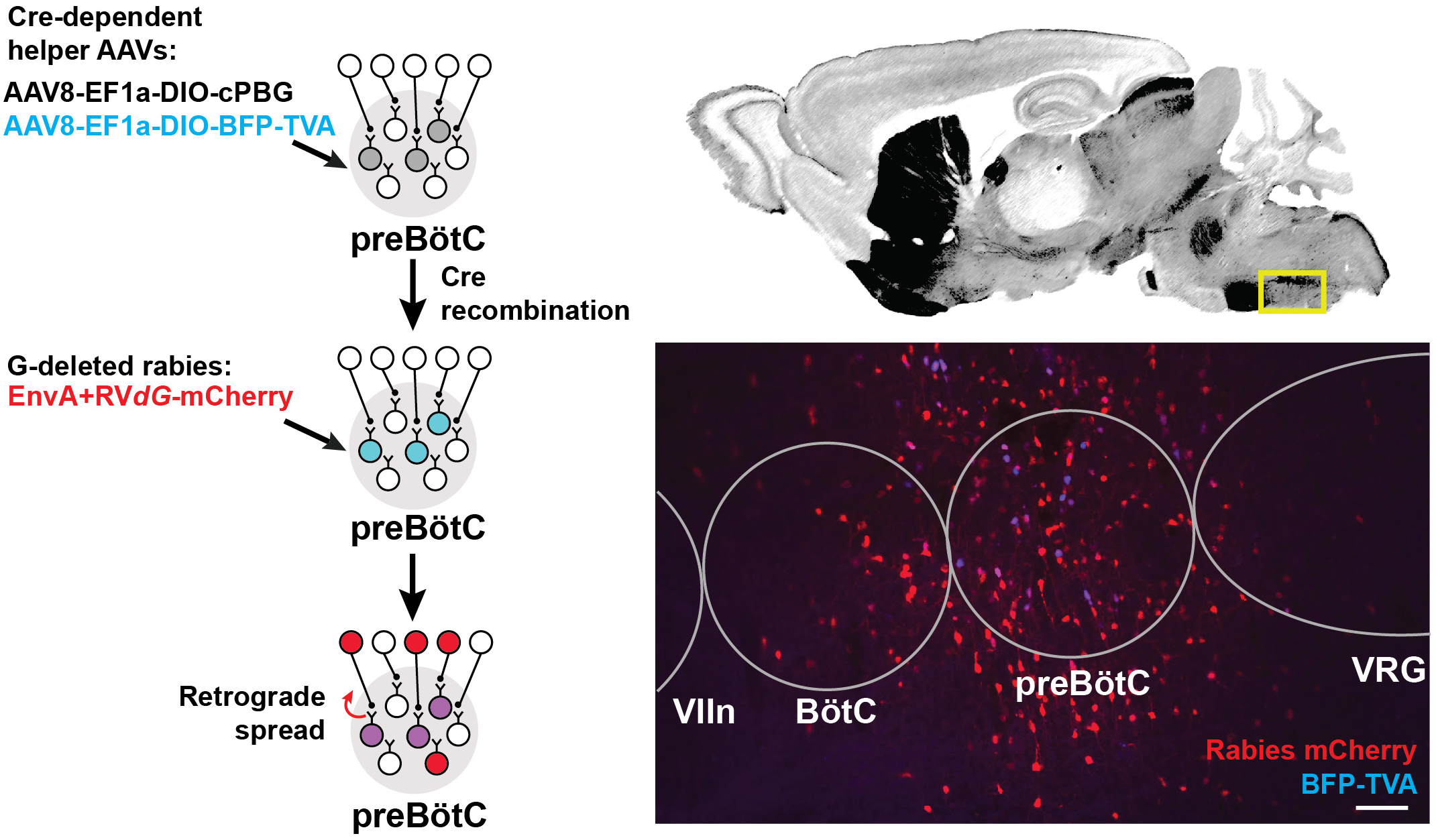
Modified rabies strategy targeting of the preBötC (left). Top right: Nissl-stained sagittal section (Paxinos & Franklin, 2004) at the level of the preBötC (yellow box). Injection of Cre-dependent helper AAVs followed by EnvA+RV*dG*-mCherry into the preBötC of SST- or GlyT2-Cre lines leads to localized expression of mCherry (red) and BFP (blue). AAV injection into GlyT2-Cre mouse leads to robust expression in the cell bodies (bottom right), allowing identification of starter cells (purple). (VIIn, facial nucleus; VRG, ventral respiratory group, rostral). Scale bar = 100 μm.

Cells infected by the rabies virus express mCherry, and co-expression of BFP from the helper AAVs enabled the identification of the putative initial Cre^+^ starter population at the target site. Neurons immediately presynaptic to the preBötC were identified as singly positive for mCherry. We note that the rabies virus may have expressed in non-Cre^+^ neurons, likely due to the leak expression of the TVA receptor, but these neurons would not be capable of long-distance, trans-synaptic rabies transfer with coincident leak expression of the glycoprotein (Kim, Juavinett, Kyubwa, Jacobs, & Callaway, 2015; Miyamichi et al., 2013; Wall, Wickersham, Cetin, De La Parra, & Callaway, 2010). Furthermore, we did not observe TVA receptor/BFP expression more than 1 mm beyond the injection site, but we do not discount the presence of lower levels of expression that we could not detect. Injection of the rabies virus alone without the helper viruses labeled <3 cells at the injection site (n=2), but not beyond. Data was discarded from experiments where there was mistargeting or extensive mCherry^+^/BFP^+^ neurons along the micropipette track or neighboring regions. We analyzed data from experiments in which mCherry^+^/BFP^+^ neurons were mostly restricted to the preBötC, with minimal double-labeled starter cells found in neighboring regions (Fig 1). Here we report the sources of afferent projections to SST^+^ and GlyT2^+^ preBötC neurons that were observed across multiple experiments (n=5 SST-Cre, n=4 GlyT2-Cre).

Given the small size of the preBötC (~500 mm across) and the inherent sparse labeling of the modified rabies strategy (Callaway & Luo, 2015), we anticipated the possibility of false negatives, i.e., inability to detect some afferent projections. To get a handle on false negatives, we used a non-genetic approach to compile a more comprehensive list of brain regions projecting to the preBötC (and its immediate surround). With this general map of afferent projections to the vicinity of the preBötC, we then carefully scrutinized these regions for neurons infected using the modified rabies strategy. We injected Fluorogold unilaterally to the preBötC and found dye uptake in many brainstem and suprapontine sites, including BötC, facial and hypoglossal nuclei, NTS, parahypoglossal region, parabrachial nuclei, Kölliker-Fuse (KF), the red nucleus, substantia nigra, motor and sensory cortex, as well as the hypothalamus and amygdala (Fig. S1).

### Afferent projections from brainstem to preBötC neurons (Figs. 2, 3)

Rabies virus injected SST-Cre mouse brains had scattered mCherry^+^ neurons in the preBötC and surround, indicating that virally infected preBötC neurons also receive synaptic input from local preBötC neurons. This spread of mCherry^+^ cells extended rostrally through the Bötzinger Complex with some neurons found in the parafacial ventral nucleus (pF_V_) and in the parafacial lateral nucleus (pF_L_) (Huckstepp, Henderson, Cardoza, & Feldman, 2016). Among dorsal regions, mCherry^+^ neurons were found in the NTS and parahypoglossal region and throughout the intermediate reticular nucleus. Some of these labeled neurons were dorsomedial to the BötC, corresponding to the region dubbed the “pre-inspiratory complex” (PiCo) (Anderson et al., 2016). Within the pons, mCherry^+^ neurons were found in the dorsolateral, medial parabrachial nuclei, KF and around the trigeminal nucleus. There were occasional neurons in the pontine reticular nuclei. In rabies-injected GlyT2-Cre mouse brains, mCherry^+^ neurons were found in all of the same regions as identified in SST-Cre injected mice though with qualitatively sparser labeling. In both SST- and GlyT2-Cre mice, mCherry^+^ neurons were visible bilaterally in any given structure, but more neurons were present on the side ipsilateral to the injection. Thus, we found that afferent projections from the same brainstem sites can innervate both excitatory SST^+^ and inhibitory GlyT2^+^ preBötC neurons.

**Figure 2.**
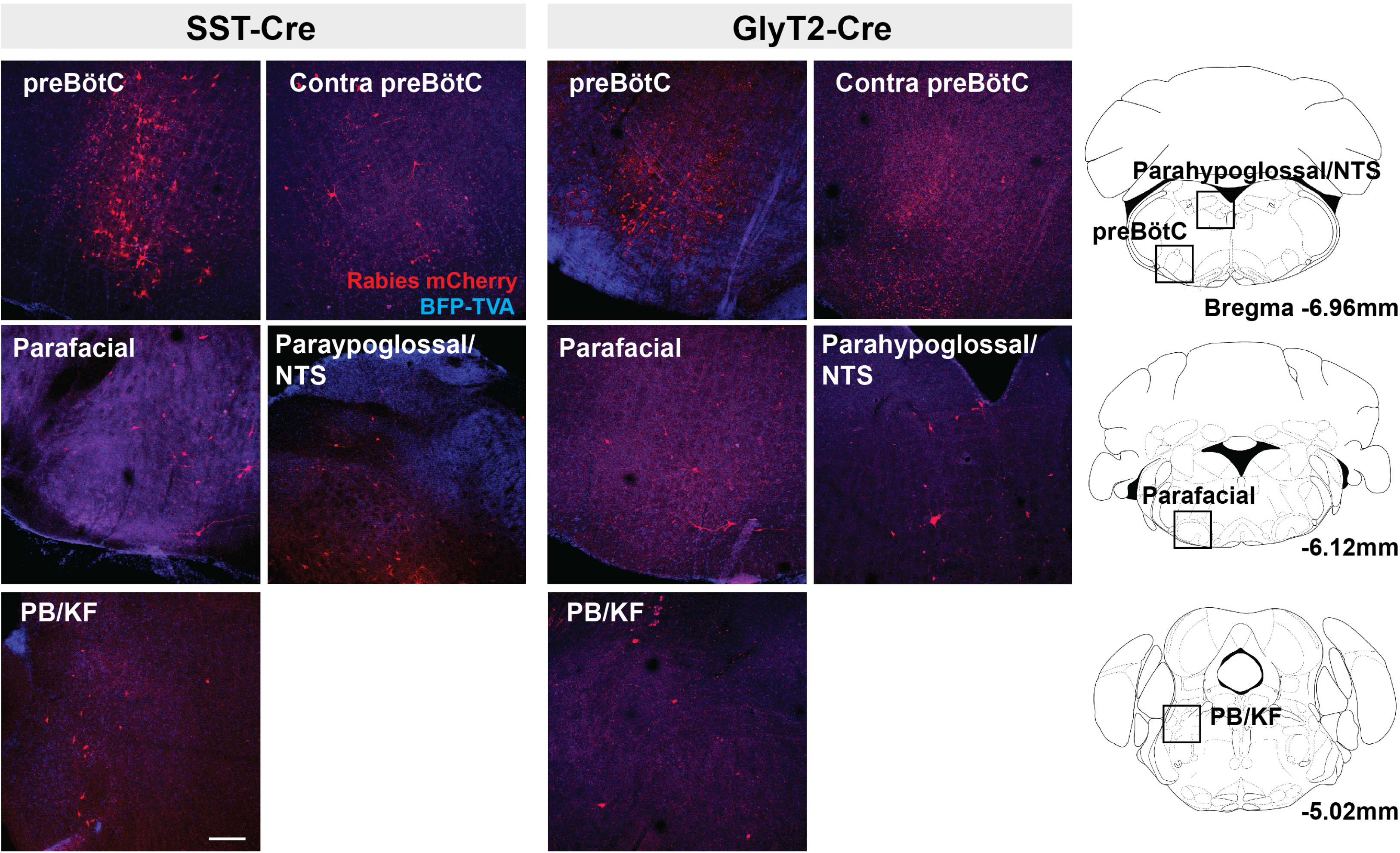
Brainstem projections of SST^+^ (left) and GlyT2^+^ (right) preBötC neurons. Left: Projections of SST^+^ neurons extend to the contralateral preBötC, the ventral respiratory group (VRG) parahypoglossal/NTS, parabrachial nuclei/KF as well as pF_V_. Right: GlyT2^+^ preBötC projections extend to the same regions. Location of each image indicated as boxes in the schematics to the right; preBötC injection site in red. Scale bar = 200μm.

**Figure 3.**
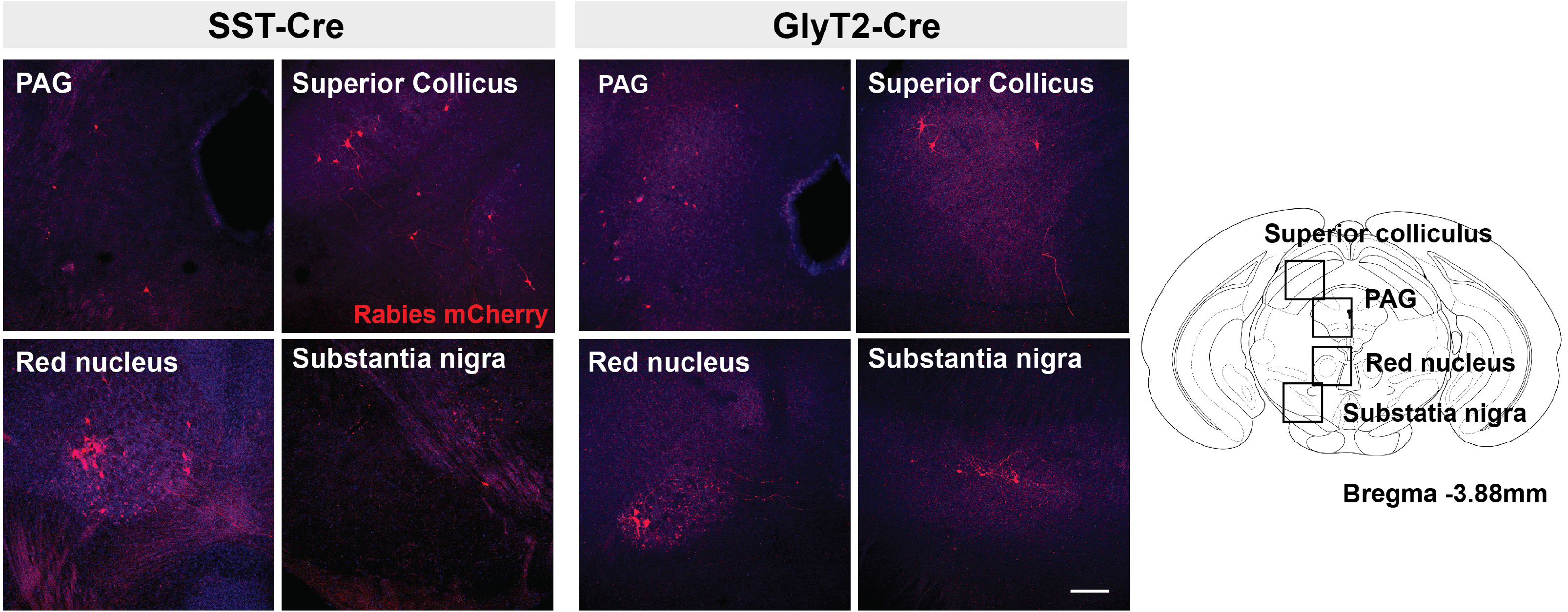
Midbrain projections of SST^+^ (left) and GlyT2^+^ (right) preBötC neurons extend to the PAG and superior colliculus, red nucleus, and substantia nigra. Boxes in schematic represent location of images shown. Scale bar = 200μm.

### Suprapontine projections to preBötC neurons (Fig. 4)

Retrogradely labeled neurons were found in all of the same midbrain regions in both SST-Cre and GlyT2-Cre mice. mCherry^+^ neurons were located bilaterally, though more neurons were found on the side contralateral to the injection. Labeled neurons were observed in the superior, but not in the inferior, colliculus, as well as in the dorsolateral PAG. In some instances (n=3 of 5 SST-Cre and n=3 of 4 GlyT2-Cre), neurons in the lateral portion of the magnocellular red nucleus were also labeled. On rare occasion (n=2 of 5 SST-Cre and n=1 of 4 GlyT2-Cre), mCherry-labeled neurons were found in the dorsal raphe nucleus and in or bordering the substantia nigra pars compacta.

**Figure 4.**
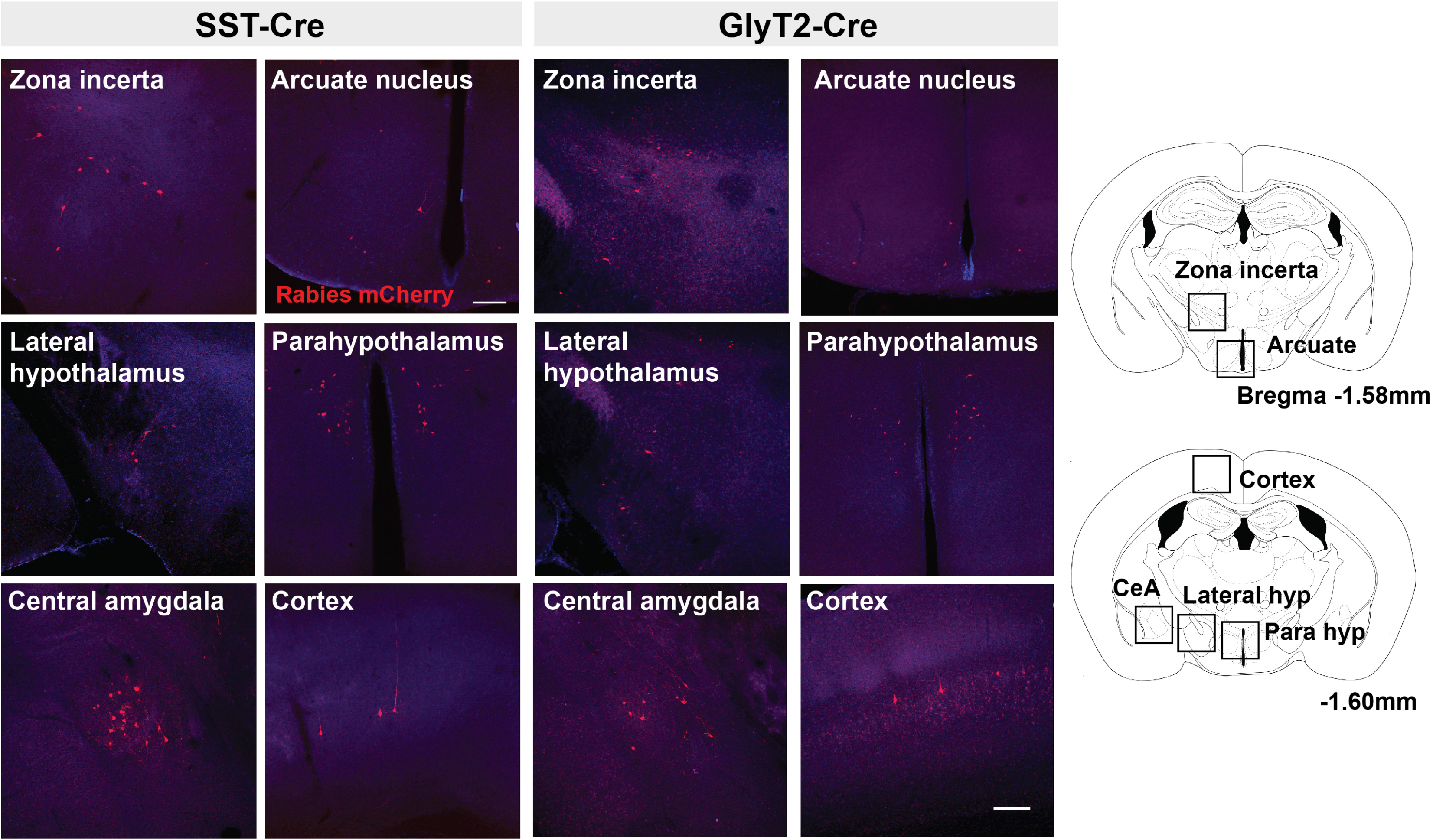
Suprapontine projections of SST^+^ (left) and GlyT2^+^ (right) preBötC neurons extend to the zona incerta, arcuate nucleus, lateral hypothalamus and parahypothalamus, as well as the central amygdala (CeA) and cortex. Boxes in schematic represent location of images shown. Scale bar = 200μm.

There were fewer mCherry-labeled neurons rostral to the midbrain, with <20 neurons in each identified region. We found mCherry^+^ neurons in the zona incerta and various compartments of the hypothalamus, including the lateral and dorsomedial hypothalamic nuclei, arcuate nucleus, and medial preoptic area. For each of the SST-Cre and GlyT2-Cre cohorts, a few retrogradely labeled neurons were found in the ventromedial and periventricular hypothalamic nuclei (n = 2 of 5 SST-Cre and 2 of 4 GlyT2-Cre), the bed nucleus of the stria terminalis (n = 2 of 5 SST-Cre and 1 of 4 GlyT2-Cre), the anteroventral periventricular nucleus of the hypothalamus, and the basal forebrain (n = 1 of 5 SST-Cre and 1 of 4 GlyT2-Cre, data not shown).

Strikingly, there were direct monosynaptic projections from the cortex and the amygdala. In cortex, mCherry^+^ neurons were found in layer L5 or L5B (as L5A neurons are cortico-cortical) of motor cortex M1. These labeled cortical neurons were scattered along the rostral-caudal axis. We found neurons in what appeared to be layers 4 and 5 of sensory cortex S1, corresponding to the hindlimb and forelimb region. Within the amygdala, retrogradely labeled neurons were sparse but consistently localized to the central compartment. For both the cortex and the amygdala, labeled neurons were present bilaterally, but more were found on the ipsilateral side of the injection. The mCherry-labeled cells were more consistently found in the amygdala and cortex of SST-Cre mice than in GlyT2-Cre mice (cortex: 3 of 5 SST-Cre mice vs 1 of 4 GlyT2-Cre mice; amygdala: 4 of 5 SST-Cre mice vs 2 of 4 GlyT2-Cre mice). Although we did not always detect mCherry^+^ neurons in every suprapontine region for each rabies-injected mouse, we did find mCherry^+^ neurons in every suprapontine region for SST-Cre as well as GlyT2-Cre cohorts, indicating that afferent neurons in each of these site project onto both excitatory and inhibitory preBötC neurons.

### Anterograde tracing from suprapontine preBötC targets (Fig. 5)

To confirm the putatively identified suprapontine afferent projections to preBötC neurons, we injected an anterograde tracer into brain regions where we observed retrogradely-labeled neurons to determine if anterograde projections from candidate regions indeed contacted preBötC neurons. Using SST reporter mice to localize the preBötC, we made large injections of tetramethylrhodamine dextran (10kMW) into: CeA, M1 motor cortex, and red nucleus. For each region injected, we found SST^+^ preBötC neurons with dextran-containing puncta. Injections into the thalamus, a region where we did not observe mCherry labeling, did not show any projections to the preBötC.

**Figure 5.**
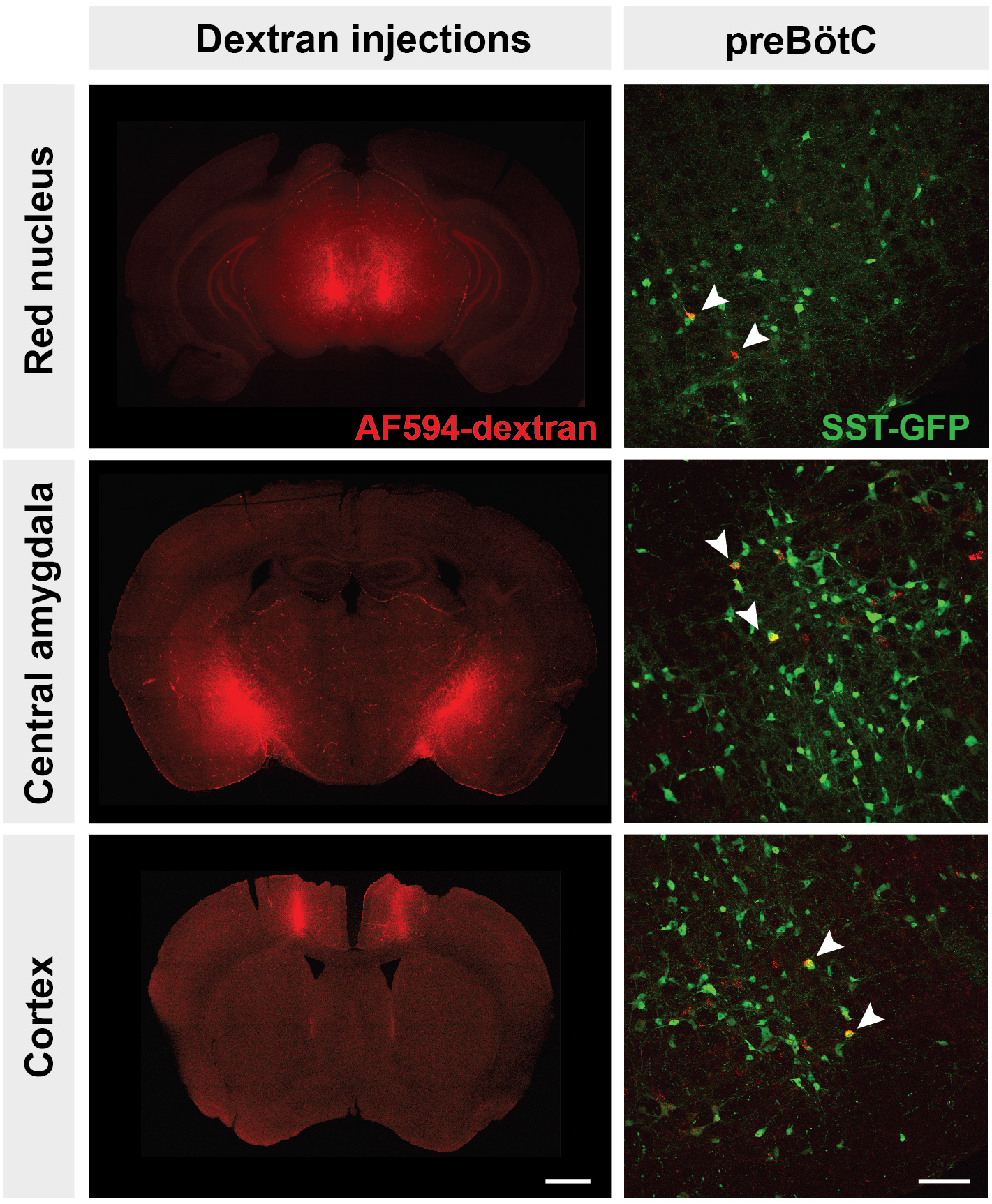
Anterograde labeling from putative suprapontine sources of afferent projections. 5% tetramethylrhodamine dextran injections into the red nucleus (top), central amygdala (middle), and cortex (bottom) (top) result in retrograde labeling of preBötC neurons. Left scale bar = 1mm; right scale bar = 100μm.

## 4. Discussion

As the kernel for the bCPG, the preBötC generates the inspiratory rhythm and transmits inspiratory-modulated signals to bulbospinal premotoneurons via projections to motoneurons innervating inspiratory muscles that underlie breathing movements. Thus, it is a ripe target for modulating breathing pattern. Here, we established multiple brainstem and suprapontine regions with monosynaptic projections to excitatory and inhibitory preBötC neurons. We first identified a panel of regions that project to the preBötC using a non-genetic tracer, allowing us to subsequently focus on regions most likely to contain retrogradely labeled cells using the modified rabies strategy. We consistently found Fluorogold-labeled neurons in regions reported to project to the preBötC but as with previous reports (Gang et al., 1995), it was impossible to limit the injection site to the preBötC or avoid transport by fibers of passage. Indeed, there were additional projections from regions that were not identified using the modified rabies strategy (see below), highlighting the specificity of our approach. Even so, given the low yield of retrograde labeling with modified rabies, we may be underestimating of the strength of these projections or failing to detect sparse projections to the preBötC neurons.

### Reciprocal preBötC projections with breathing-related regions in the brainstem

Many brainstem regions reciprocally connect with the preBötC, representing recurrent circuits for feedback modulation of breathing. The bCPG includes a column of respiratory-related nuclei along the ventral medulla that interact with the preBötC (Alheid & McCrimmon, 2008). We now show that many of these regions project directly to both excitatory SST^+^ and inhibitory GlyT2^+^ preBötC neurons. The BötC, just rostral to the preBötC, contains mostly inhibitory glycinergic neurons that are active during postinspiration and expiration (Ezure, 1990; Ezure & Manabe, 1988; Ezure, Tanaka, & Kondo, 2003; Jiang & Lipski, 1990; Tian, Peever, & Duffin, 1999) that receives projections from parabrachial/KF nuclei, NTS, and the parafacial region (Gang et al., 1995). Within the pF, we find more projections from the pF_V_ to the preBötC than the pF_L_. As the p_FV_ provides chemosensory-related excitatory tone for breathing, it can do so via direct projections to preBötC SST^+^ and GlyT2^+^ neurons, in addition to downstream projections to hypoglossal and phrenic (pre)motoneurons (Chamberlin, Eikermann, Fassbender, White, & Malhotra, 2007; Rosin, Chang, & Guyenet, 2006). There are projections originating dorsomedial to pF and BötC, the site of the presumptive PiCo (Anderson et al., 2016) to the preBötC; this projection could act to coordinate and sequence inspiration and post-inspiration.

The preBötC receives descending and ascending signals from other brainstem regions that can modulate its activity to affect breathing. The NTS receives sensory signals from vagus, facial, hypoglossal and glossopharyngeal nerves, as well as peripheral chemo-, baro-, and mechano-receptors; it also receives descending input from the forebrain originating from hypothalamus, including the paraventricular nucleus, amygdala (basolateral, central), and cortex (Affleck, Coote, & Pyner, 2012; Schwaber, Kapp, Higgins, & Rapp, 1982; van der Kooy, Koda, McGinty, Gerfen, & Bloom, 1984) that can modulate the processing of these sensory signals. The NTS also contains premotoneurons critical for coordinating expiratory breaths for vocalizations (Hernandez-Miranda et al., 2017). From the pons, the preBötC receives direct input from parabrachial and KF nuclei, which play a role in regulating the timing and phase transitions of the respiratory cycle. These regions are implicated in coupling vocalization, swallowing, and nociception to breathing (Dutschmann & Dick, 2012). The dorsolateral pons receives projections from the NTS, pF_V_, amygdala, bed nucleus of the stria terminalis, hypothalamus, substantia nigra, and ventral tegmental area, thus providing the preBötC with integrated input from many modalities.

### Suprapontine projections to preBötC relay integrated sensorimotor information

Projections to the preBötC from the midbrain originate from structures that receive sensory information and process motor commands, therefore providing a means by which breathing could be modulated and coordinated with other behaviors. The periaqueductal gray (PAG) is postulated to integrate motor, limbic, and sensory information to modulate breathing across behaviors, such as gasping and vocalization, in a context-specific manner (Subramanian & Holstege, 2010). Stimulation of the PAG can change preBötC activity, and activation of different PAG divisions results in distinct breathing patterns. The superior colliculus transforms sensory information into motor output or behaviors, e.g., orienting, approach, or defensive behaviors. Stimulation of the superior colliculus can change breathing. With direct projections to the preBötC from the superior colliculus, this pathway could coordinate breathing with specific behaviors, with minimal delay. While both the superior and inferior colliculi receive efferent projections from preBötC, retrogradely labeled neurons were almost exclusively in the superior colliculus. As the inferior colliculus primarily processes auditory signals, this suggests that other, perhaps indirect, pathways may coordinate auditory-related behaviors with breathing or underlie audition-induced changes in breathing, as in the startle response (Fendt, Li, & Yeomans, 2001).

Both the substantia nigra and the red nucleus participate in motor coordination, but neither has been directly linked to neural pathways affecting breathing. An indirect pathway between the substantia nigra and the retrotrapezoid nucleus through the PAG is impacted in a Parkinson’s model of respiratory failure in rats (Lima, Oliveira, Botelho, Moreira, & Takakura, 2018), and red nucleus neurons may play a role in regulating ventilatory response to hypoxia (Ackland, Noble, & Hanson, 1997). As the red nucleus and the substantia nigra are responsible for coordinating and facilitating movement, these regions could be relaying information to the preBötC to synchronize breathing with gait or other movements.

Descending afferent projections from the forebrain provide information regarding emotional, cognitive, and physiological state to the preBötC. With various hypothalamic nuclei projecting to the preBötC, there are potentially a broad range of regulatory behaviors that could directly and rapidly influence breathing. The arcuate regulates feeding behavior, energy homeostasis, and diurnal temperature, while the medial preoptic area is implicated in temperature regulation and sex-specific behaviors such as mating and aggression. The lateral and dorsomedial hypothalamus are involved in feeding, reward, sleep/wakefulness, and stress. These regions are rich in neurons that secrete peptides and hormones such as ghrelin, orexin, and melanin-concentrating hormone (MCH), which can affect the breathing across different arousal states (Benedetto et al., 2013). The hypothalamus is highly interconnected with the limbic and motor systems and could provide information to modulate breathing in relation to physiological or behavioral state.

Projections from the cortex to the preBötC were present throughout primary and secondary motor cortex. Although corticobulbar neurons from primary motor cortex innervate premotor and motoneurons in and around the trigeminal, facial, vagus, hypoglossal, and glossopharyngeal nerves, no direct connections from the cortex to the preBötC had been identified. These projections could provide efference copy to the preBötC for behaviors that need to be timed with the breathing cycle, e.g., chewing, swallowing, and vocalization (Martin-Harris, 2006; Rea, 2015).

The CeA is the output pathway from amygdala to brainstem. While it contributes to the learning and expression of fear responses, e.g., fight or flight and emotional behavior, the CeA also modulates physiological responses to stress. Direct stimulation of the CeA across the breathing cycle results in a shift to inspiratory phase, and the activity of some CeA neurons are phase-locked with the breathing cycle (Frysinger, Zhang, & Harper, 1988; Harper, Frysinger, Trelease, & Marks, 1984). In humans, electrical stimulation of the amygdala results in an apnea (Dlouhy et al., 2015), suggesting that the amygdala is functionally coupled to the brainstem bCPG. The CeA receives input from cortex, limbic system, hypothalamus, olfactory pathway, substantia nigra, and many brainstem nuclei, and processes information to generate a breathing response to emotional stimuli, mediated at least in part by direct projections to the bCPG.

### Functional input to preBötC neurons

Afferent inputs to the preBötC originate from different regions of the brain associated with different functions and contain neurons with diverse molecular phenotypes that will also likely corelease peptidergic and/or hormonal neuromodulators that influence breathing pattern, e.g., opioids, serotonin, orexin, SST, ghrelin, and MCH (Gray, Janczewski, Mellen, McCrimmon, & Feldman, 2001; Mulkey et al., 2007; Young et al., 2005). Depending on the phenotype of these neurons, the onset and duration of effects on breathing could span a broad range. Since afferent projections from each region project to both SST^+^ and GlyT2^+^ preBötC neurons, the net effect on preBötC activity will depend on whether the inputs from individual neurons are targeting one or both subpopulation(s), as well as their neurotransmitter/neuromodulator release and the associated postsynaptic receptors. Indeed, stimulation within the amygdala, even just the CeA, can result in a multitude of different breathing responses (Bonvallet & Gary-Bobo, 1971; Dlouhy et al., 2015).

### Reciprocal projections to the preBötC allow feedback control

SST^+^ and GlyT2^+^ preBötC neurons project broadly within the brainstem and have select efferent suprapontine targets (Yang & Feldman, 2018). Many brainstem regions have reciprocal projections with the preBötC including pF, BötC, NTS, and parabrachial/KF nuclei that could modulate and/or regulate the precise pattern of breathing. Surprisingly many suprapontine regions including superior colliculus, dorsomedial and lateral hypothalamus, and zona incerta also have reciprocal projections with excitatory and inhibitory preBötC neurons. Other regions projecting to preBötC, e.g., cortex, CeA, inferior colliculus, that do not appear to receive direct projections from SST^+^ or GlyT2^+^ preBötC neurons, could receive inputs from other preBötC cell types, e.g., SST^−^ glutamatergic or GABAergic neurons. Moreover, these regions could receive oligosynaptic input arising from preBötC. Respiratory activity can be detected in and phase-locked with oscillations in the cortex, hippocampus, and amygdala, (Rojas-Libano, Wimmer Del Solar, Aguilar-Rivera, Montefusco-Siegmund, & Maldonado, 2018), and such entrainment is postulated to contribute to cognitive functions including memory encoding and consolidation. In addition to ascending respiratory-related oscillations that appear to originate in olfactory sensory neurons, direct or indirect pathways from bCPG are also capable of relaying respiratory rhythms to the forebrain. Although we did not detect direct projections from hippocampus, prefrontal cortex, orbital cortex, or olfactory bulb to the brainstem, we did find such projections from motor cortex, central amygdala, and hypothalamic nuclei. Essential to note is that brain regions with sparse projections to the preBötC could be missed with our assay. Nonetheless, the direct afferent projections from the forebrain to the preBötC that we identified can represent a substrate for coordinating inspiratory activity with other behaviors or physiology, e.g., cardiorespiratory coupling, vocalizations, whisking, feeding, sniffing, or cognitive processes.

### Direct/indirect control of breathing

The preBötC neuronal subpopulations targeted for retrograde tracing were either SST^+^ or GlyT2^+^, which accounts for a fraction of its excitatory and inhibitory population (~17% and ~50%, respectively). Despite such a small target population, we identified a select list of brainstem and suprapontine regions that project directly onto preBötC neurons, indicating that the bCPG receives information across many modalities and suggesting considerable and complex modulation by emotional- and cognitive-related input. In addition to the input onto these preBötC populations, afferent information may also be introduced into the bCPG via targeting rhythmogenic preBötC neurons. Delineating the afferent input into the preBötC thus captures the flow of information in generating the inspiratory motor command and reveals pathways for higher-order control of breathing.

## Acknowledgements

The authors would like to thank Grace Li for excellent technical assistance and the Feldman lab for comments on the manuscript. This work was supported by the A.P. Giannini Foundation (CFY) and the National Institutes of Health (NIH F32 HL126522 (CFY), RO1 EY022577 and MH063912 (EMC), RO1 NS72211 and R35 HL135779 (JLF)).

---

Figure S1. Retrograde labeling of putative afferent projections to the preBötC. Fluorogold injections into the preBötC (top, arrow) result in retrograde labeling of neurons throughout the brain including the nucleus of the solitary tract (NTS) contralateral preBötC, parafacial nucleus, parabrachial nuclei and Kölliker-Fuse (PB/KF), trigeminal nucleus (MO5), substantia nigra, red nucleus, superior colliculus, central amygdala, lateral hypothalamus and zona incerta, parahypothalamus, bed nucleus of the stria terminalis (BNST), lateral preoptic area (LPO), and cortex. Top scale bar = 500um, bottom scale bar = 200μm.

