## Supplementary figures and images for "Monosynaptic projections to excitatory and inhibitory preBötzinger Complex neurons"

### Fig S1

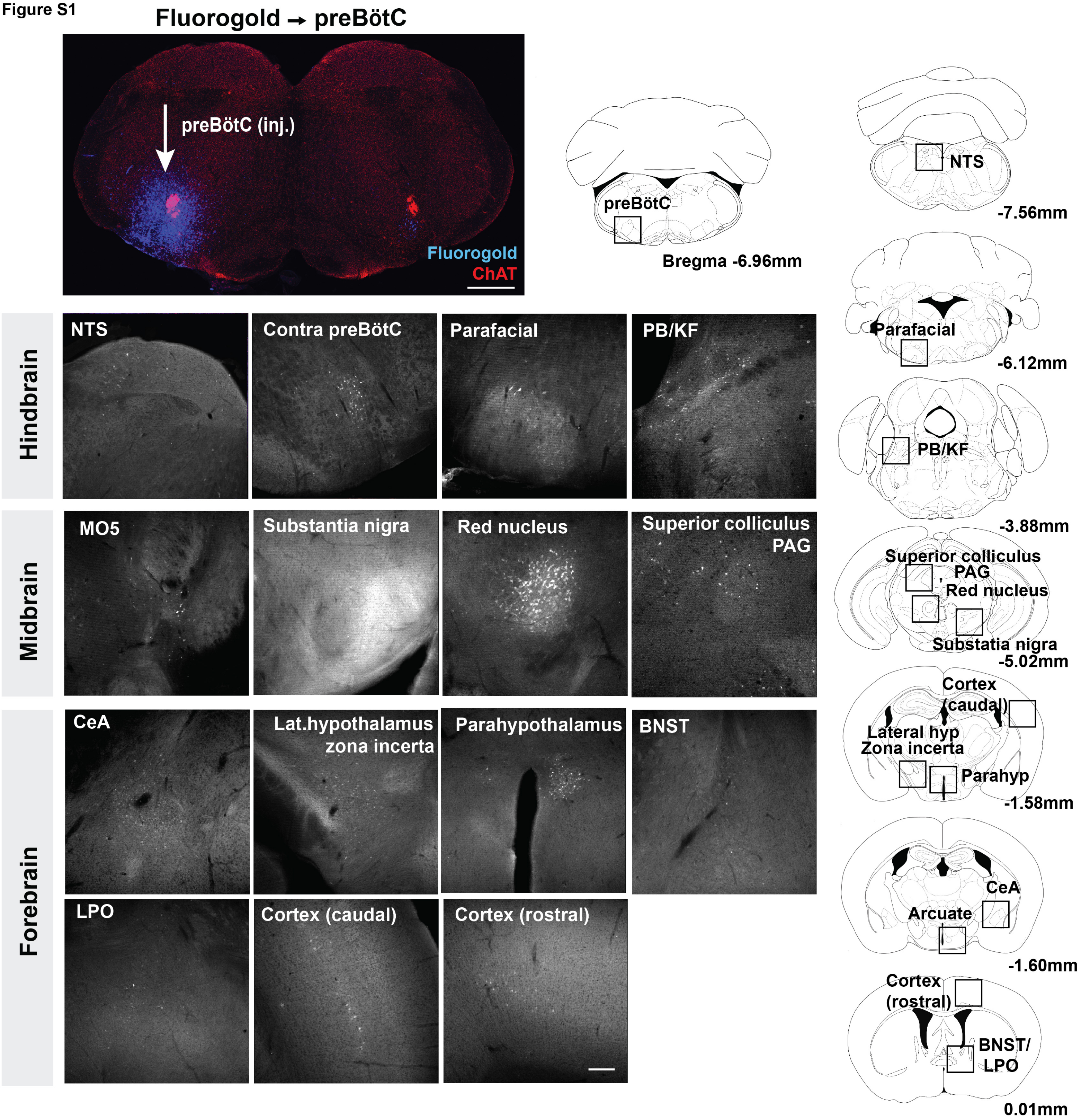
